# Reciprocal BLUP: A Predictability-Guided Multi-Omics Framework for Plant Phenotype Prediction

**DOI:** 10.1101/2025.11.27.690927

**Authors:** Hayato Yoshioka, Gota Morota, Hiroyoshi Iwata

## Abstract

Sustainable improvement of crop performance requires integrative approaches that link genomic variation to phenotypic expression through intermediate molecular layers. Here, we present Reciprocal Best Linear Unbiased Prediction (Reciprocal BLUP), a predictability-guided multi-omics framework that quantifies cross-layer relationships among the genome, metabolome, and microbiome to enhance phenotype prediction. Using a panel of 198 soybean accessions grown under well-watered and drought conditions, we first evaluated four direction-specific prediction models (genome→microbiome, genome→metabolome, metabolome→microbiome, and microbiome→metabolome) to estimate the predictability of individual omics features. Then, we evaluated whether subsets of features with high cross-omics predictability improve phenotype prediction. These cross-layer models identify features that play physiologically meaningful roles within multi-omics systems, enabling us to prioritize variables that capture coherent biological signals enriched for phenotype-relevant information. As a result, metabolome features were highly predictable from microbiome data, whereas microbiome predictability from metabolomic data was weaker and more environment-dependent, revealing an asymmetric relationship between these layers. In the subsequent phenotype prediction analysis, the model incorporating predictability-based feature selection substantially outperformed models using randomly selected features and also achieved prediction accuracies comparable to those of the full-feature model. Under drought, phenotype prediction models based on metabolomic or microbiomic kernels (MetBLUP or MicroBLUP) outperformed the genomic baseline (GBLUP) for several biomass-related traits, indicating that environment-responsive omics layers captured phenotypic variation not explained by additive genetic effects. Our results highlight hierarchical interactions among genomic, metabolic, and microbial systems, with the metabolome functioning as an integrative mediator linking genotype, environment, and microbiome composition. The Reciprocal BLUP framework provides a biologically interpretable and practical approach for integrating multi-omics data, improving phenotype prediction, and guiding omics-based feature selection in plant breeding.

## 1. Introduction

Sustainable increases in agricultural productivity are essential to meet the demands of a growing global population. To address these challenges, modern plant breeding increasingly leverages genomic prediction approaches such as Genomic Best Linear Unbiased Prediction (GBLUP), which enable accurate estimation of breeding values using genome-wide marker information [1]. While genomic information alone can capture substantial genetic variance, complex traits are also shaped by intermediate molecular and ecological processes that are not fully explained by additive genetic effects [2,3].

Recent advances in multi-omics technologies have made it possible to profile diverse biological layers—genome [4], metabolome [5], and microbiome [6]—within the same individuals. These intermediate molecular layers provide unprecedented opportunities to unravel the complex causal relationships among biological systems, as they can act as mediators linking genotypes to phenotypes [7,8].

Among them, rhizosphere microorganisms play a pivotal role in plant growth and resilience by enhancing nutrient uptake, suppressing pathogens, and improving tolerance to abiotic stresses such as drought [9–11]. Understanding the structure and function of the rhizosphere microbiome is therefore essential for deciphering the mechanisms of plant–microbe symbiosis and their contributions to stress adaptation [12].

In addition, the metabolome provides a complementary dimension by capturing the dynamic physiological state of plants under varying environmental conditions. Because metabolite levels integrate genetic, environmental, and microbial influences, they can serve as sensitive biomarkers of stress responses and genotype-by-environment interactions [13,14].

Despite these advances, most existing frameworks treat all omics features equally, potentially diluting biologically meaningful signals. While Christensen et al. [15] and Zhao et al. [16] focused on how to handle missing intermediate omics information within genetic evaluation frameworks, the distinct question of feature selection—identifying which omics variables are biologically relevant and predictive—has been less explored.

The need for feature selection arises in part from the characteristics of intermediate omics layers. These layers are high-dimensional and often contain substantial redundancy and noise, making it challenging to determine which features truly contribute to transmitting biologically meaningful signals toward the phenotype. This motivates the need for biologically informed strategies to prioritize omics features.

In this study, we aim to develop a reciprocal multi-omics framework that leverages predictability among genomic, metabolomic, and microbiomic layers to guide feature selection. We quantify interactions and variance components across these interconnected layers in a plant–microbe system, and evaluate whether cross-omics–informed feature subsets enhance phenotype prediction under contrasting environmental conditions.

To support this framework, we introduce the concept of cross-omics predictability—the extent to which one omics layer can be predicted from another—as a biologically grounded criterion for feature prioritization. This metric highlights features that occupy functionally meaningful positions within the information flow from genotype to phenotype. Highly predictable features tend to represent coherent biological variation, including upstream–downstream regulatory relationships, stable genetic or ecological influences, and consistent responses to environmental perturbations. Therefore, prioritizing such features provides a principled strategy for dimensionality reduction that enriches for phenotype-relevant signals while mitigating noise, potentially improving phenotype prediction accuracy.

In the following sections, we first describe the multi-omics datasets and experimental design employed in this study (Section 2.1). We then introduce our predictability-based integration framework, outlining the implementation of inter-omics prediction and the identification of predictable features (Section 2.2), and describe how these selected omics features were used for phenotype prediction relative to genomic and random baselines (Section 2.4). The results of these analyses are presented in Section 3, followed by a discussion of their biological implications for understanding plant–microbe–environment interactions and improving genomic prediction frameworks (Section 4).

## 2. Materials and Methods

### 2.1. Soybean Multi-Omics Data

We analyzed multi-omics datasets collected from a common panel of *N* = 198 soybean accessions. The panel was drawn from the Global Soybean Minicore Collection [4]. The datasets included:

- **Whole-genome genotypes (genome)**: a 198 *×* 425,858 matrix, where each column corresponds to a single nucleotide polymorphism marker in the soybean genome [4].
- **Rhizosphere metabolome**: a 198 *×* 265 matrix of metabolome features obtained from rhizosphere samples. Each column represents the normalized peak area of a distinct metabolome feature detected by tandem mass spectrometry [17].
- **Rhizosphere microbiota profiles (microbiome)**: a 198 *×* 16,457 matrix generated by 16S rRNA gene amplicon sequencing of DNA extracted from rhizosphere samples. Each column represents the relative abundance of an amplicon sequence variant (ASV) [17].
- **Plant phenotypes**: a 198 *×* 9 matrix. These traits include biomass-related features such as shoot and leaf fresh/dry weights, growth stage, plant height, stem length, number of nodes, and number of branches.

All measurements were obtained from field trials conducted in 2019 under two watering conditions: a well-watered regime (control) and a water-limited regime (drought), enabling assessment of environmental effects. Microbiome and metabolome datasets are identical to those described in Dang et al. [17], and were collected from the same field plots as the phenotypic measurements to ensure sample consistency across omics layers. Microbiome data were filtered to remove chloroplast and mitochondrial sequences, and all datasets were subsequently scaled prior to downstream analyses. The details of the data collection are described in Appendix A.

### 2.2. Predictability analysis among omics layers

We implemented a Best Linear Unbiased Predictor (BLUP)-based framework to integrate multiple omics layers through predictability-guided feature selection and phenotype prediction. Although BLUP was originally formulated to predict random effects based on expected pedigree relationships, its linear mixed-model framework was later extended to incorporate genome-wide markers (GBLUP). This extension enables efficient analysis of high-dimensional genomic data [1].

This framework quantifies how omics features represent genetic and environmental signals, identifies stable high-predictability features across environments, and evaluates their contribution to the prediction of phenotypic values. The analysis focuses on interactions among genome, microbiome, and metabolome layers under both control and drought conditions.

To characterize how different omics layers capture genetic and environmental information, we performed four direction-specific prediction analyses. We employed BLUP models, each fitted with a single omics-derived relationship matrix (Fig. 1):

**Figure 1.**
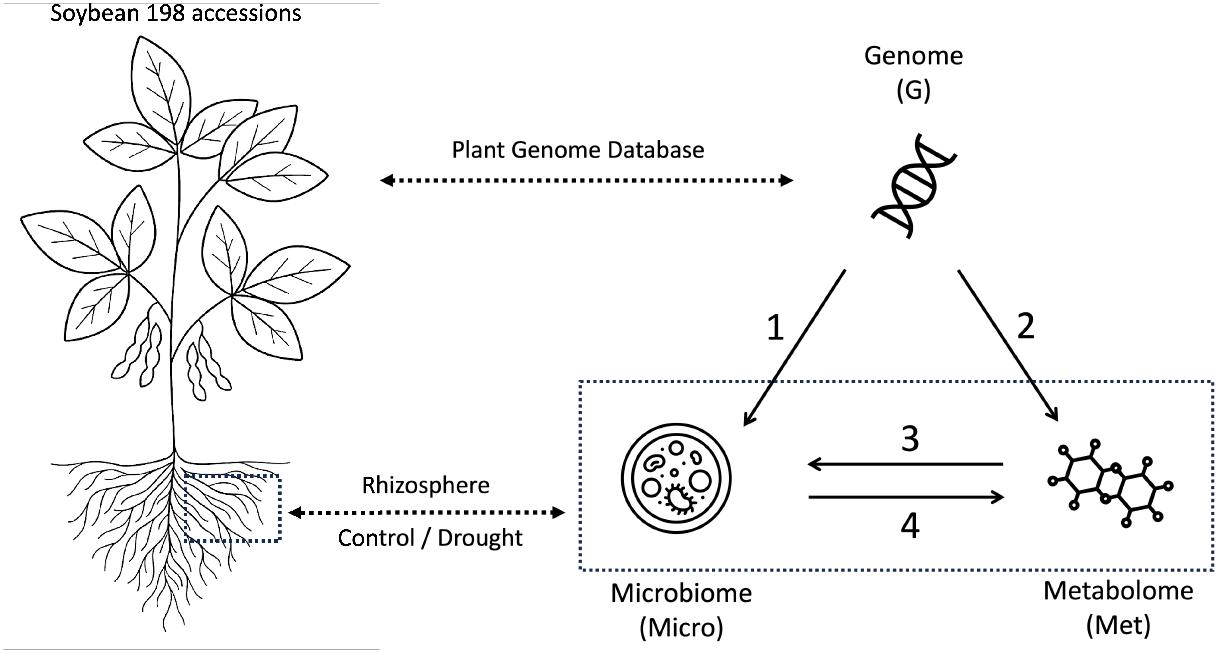
Conceptual overview of the modelling framework. We investigated how genomic (G), microbiome (Micro), and metabolome (Met) layers contribute to phenotype prediction across environments. Four direction-specific BLUP models were considered: (1) genome → microbiome, (2) genome → metabolome, (3) metabolome → microbiome, and (4) microbiome → metabolome. The microbiome and metabolome were sampled from the rhizosphere. The omics features were ranked by predictability in these directions, and the selected top features were evaluated in phenotype prediction.

**Model 1:** genome *→* microbiome

**Model 2:** genome *→* metabolome

**Model 3:** metabolome *→* microbiome

**Model 4:** microbiome *→* metabolome

Let **M** = (**m**_1_, …, **m**_*q*_) denote the omics matrix to be predicted (e.g., metabolome or microbiome), and **X** the explanatory omics used as predictors. Here, when **M** represents the metabolome, **m**_*j*_ denotes the vector of observed values for the *j*-th metabolite; whereas when **M** represents the microbiome, **m**_*j*_ denotes the vector of observed abundances for the *j*-th ASV. For each response vector **m**_*j*_, we fit a BLUP model with a single omics-derived relationship matrix:

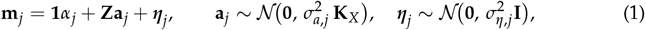

where *α*_*j*_ is the fixed-effect intercept for feature **m**_*j*_, **Z** is the incidence matrix linking samples to the random effects, **a**_*j*_ represents the vector of omics-based random effects, and ***η***_*j*_ is the residual error term. The variance component 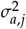 denotes the variance of the omics-based random effects, whereas 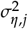 represents the residual variance, both specific to feature **m**_*j*_.

The matrix **K**_*X*_ represents the covariance structure of the omics-derived random effects. It is constructed from the explanatory omics data **X** as a linear relationship matrix:

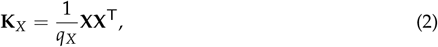

where *q*_*X*_ denotes the number of explanatory features.

Predictive performance was evaluated by 5-fold cross-validation, and quantified by Pearson’s correlation (COR) between observed and predicted values:

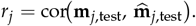

### 2.3. Feature selection based on predictability

Features were ranked according to their predictability score *r*_*j*_, which quantifies how well each omics feature can be predicted from other data sources. Instead of applying a fixed correlation threshold, we selected features based on their relative predictability ranking. Specifically, we considered the top 100%, 50%, 25%, 12.5%, and 6.25% of features, corresponding to progressively stricter levels of selection.

For each environment *e ∈ {control, drought}*, the top predictable features were defined as the top *p*% of features ranked by 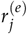:

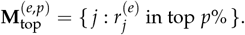

Here, *p* denotes the proportion of features retained.

Because a given omics layer may be predictable from multiple information sources, we combined results across relevant models. For the metabolome, three feature selection strategies were applied:

1. **top:G** — metabolites highly predictable from the genome;
2. **top:Micro** — metabolites highly predictable from the microbiome;
3. **top:G+Micro** — metabolites highly predictable from genome and microbiome. Similarly, for the microbiome, features were selected under three analogous rules:

1. **top:G** — microbes predictable from the genome;
2. **top:Met** — microbes predictable from the metabolome;
3. **top:G+Met** — microbes predictable from genome and metabolome.

These top-feature sets 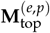 were used for downstream phenotype prediction analysis.

### 2.4. Phenotype prediction using selected omics features

To evaluate the contribution of predictable omics features to phenotype prediction, we constructed linear kernels—used here as sample-to-sample covariance structure (relationship matrices)—from the features in each omics layer as follows:

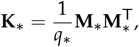

where **M**_*∗*_ represents the feature matrix used to compute the kernel and *q*_*∗*_ is the number of features included. We compared three feature sets:

- *Full omics* 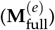 : all available features in the omics layer were used to construct the kernel, representing the baseline performance without feature filtering.
- *Top omics* 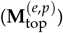: only the top *p*% of features ranked by predictability in environment *e* were used. These represent the biologically and statistically predictable subset identified through the predictability analysis.
- *Random omics* 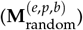: a random subset of features with the same size as 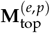, generated for each iteration index *b* (*b* = 1, …, 10) to benchmark random selection against predictability-based selection.

These three settings enabled us to disentangle the effects of feature selection and model complexity on phenotype prediction accuracy.

Phenotype prediction was then performed using a BLUP model:

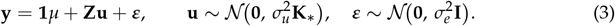

Here, **y** denotes the vector of phenotypic values, *μ* is the fixed-effect intercept, and **Z** is the design matrix from samples to random effects. The vector **u** represents random effects with variance component 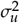, whereas ***ε*** denotes residual errors with variance 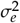.

Models based on metabolome and microbiome kernels are denoted as *MetBLUP* and *MicroBLUP*, respectively. The GBLUP model using the additive genomic relationship matrix **K**_G_, calculated according to Eq. (2), was included as a baseline.

To account for variability in random feature selection, the entire 5-fold cross-validation procedure (using five different seeds) was repeated 10 times, with a new random subset 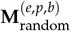 generated at each iteration (*b* = 1, …, 10).

All analyses were implemented in R [18] using the RAINBOWR package [19], and visualizations were produced with ggplot2 [20].

## 3. Results

### 3.1. Predictability patterns across omics layers and environments

Pairwise prediction performance across omics layers is summarized in Fig. 2. Microbiome prediction was more accurate when metabolome features were used as predictors (Model 3: Met*→*Micro) than when genome features were used as predictors (Model 1: G*→*Micro), under both control and drought conditions. Conversely, the prediction of the metabolome was more accurate when microbiome features served as predictors, particularly under drought conditions (Model 4: Micro*→*Met).

**Figure 2.**
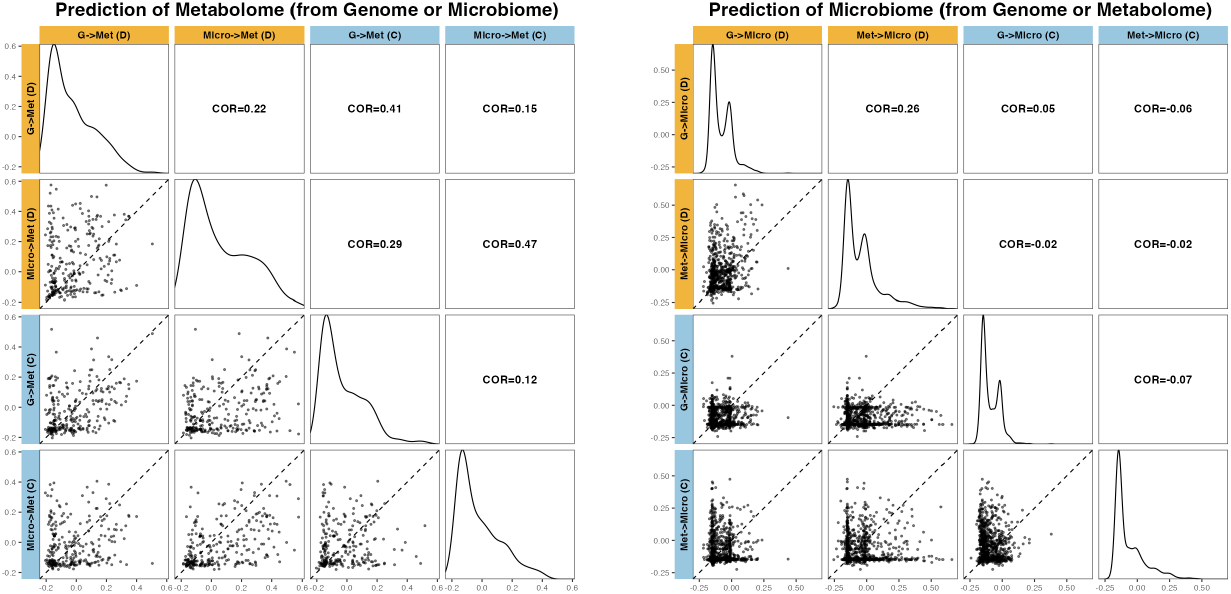
Prediction accuracy for each omics by method and environment. Left: (a) Metabolome; Right: (b) Microbiome. For the microbiome, Model 1 (G → Micro) and Model 3 (Met → Micro); for the metabolome, Model 2 (G → Met) and Model 4 (Micro → Met) are compared under Drought (D) and Control (C) conditions. Diagonal panels show the distributions of prediction accuracy of each omics feature for each setting; lower triangles display pairwise scatter plots of prediction accuracy for each omic feature (the dashed line indicates *y*=*x*); upper triangles report correlation coefficients between settings.

The similar pattern was observed in the number of features selected based on predictability thresholds (Table 1). Model 1 (G*→*Micro) selected far fewer features with *r >* 0.1, only 10/1193 (0.8%) under control and 35/893 (3.9%) under drought conditions. In contrast, Model 3 (Met*→*Micro) selected a larger number of metabolome features with high predictability (*r >* 0.1), identifying 132/1193 (11.1%) under control and 127/893 (14.2%) under drought conditions.

**Table 1.** Number of selected features. The correlation (r) threshold for each direction-specific prediction model. The *Full* column shows the total number of available features without correlation filtering.

| Model | Environment | $r > 0.3$ | $r > 0.2$ | $r > 0.1$ | $r > 0$ | Full |
| --- | --- | --- | --- | --- | --- | --- |
| Model 1: G → Micro | Control | 1 | 3 | 10 | 91 | 1193 |
|  | Drought | 1 | 4 | 35 | 105 | 893 |
| Model 2: G → Met | Control | 8 | 16 | 55 | 95 | 265 |
|  | Drought | 7 | 28 | 58 | 91 | 265 |
| Model 3: Met → Micro | Control | 15 | 55 | 132 | 263 | 1193 |
|  | Drought | 38 | 68 | 127 | 233 | 893 |
| Model 4: Micro → Met | Control | 11 | 22 | 56 | 95 | 265 |
|  | Drought | 39 | 72 | 98 | 131 | 265 |

Similarly, Model 4 (Micro*→*Met) under drought conditions identified more predictable metabolome features than under control conditions, suggesting enhanced metabolic signaling to microbial structure in water-limited environments.

### 3.2. Robust metabolome features across conditions and models

A Venn diagram summarizing features exceeding *r >* 0.1 across the four models and two environments (Fig. 3) revealed a set of robust metabolome features consistently predictable regardless of model direction or environment. Three metabolites—glutamic acid, daidzein, and genistin—were consistently selected across all conditions. These compounds have been implicated in root exudation, plant–microbe interactions, and stress responses in the rhizosphere, underscoring their biological relevance [21–23].

**Figure 3.**
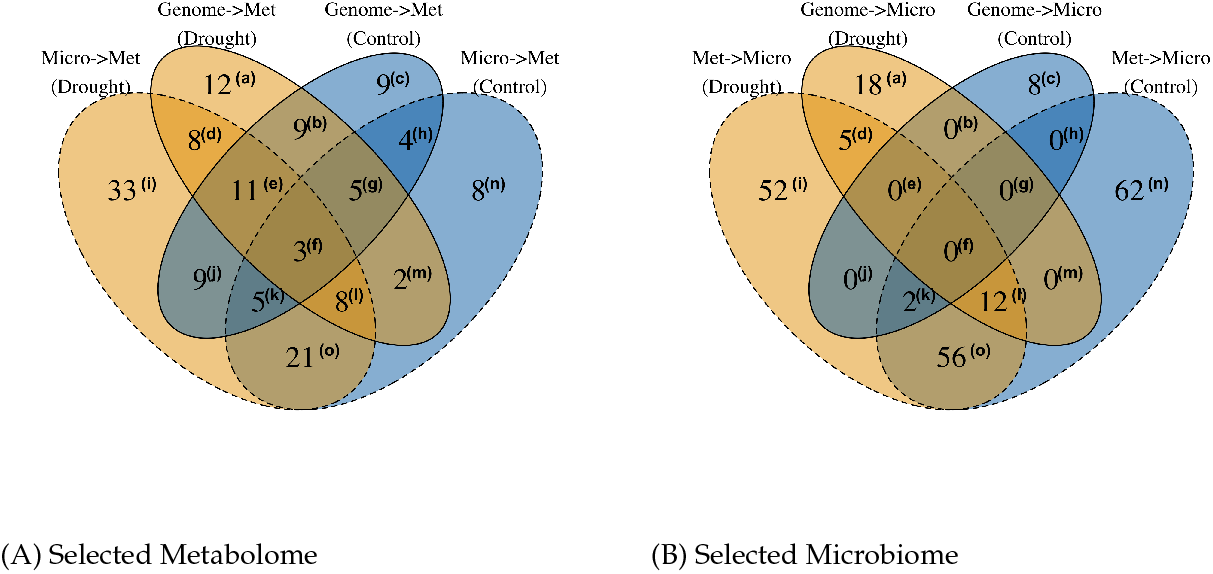
The selected microbiome and metabolome features under each condition. Drought: yellow; Control: blue. Identified using Pearson correlation (*r >* 0.1). (A) For the prediction of the metabolome, genomic or microbiome features were used as inputs. (B) For the prediction of the microbiome, genomic or metabolome features were used as inputs.

In contrast, no single bacterial taxon was consistently predictable across environments or models. Microbiome feature selection exhibited pronounced environment-and genome-dependent variation.

### 3.3. Phenotype prediction using predictability-selected omics features

#### 3.3.1. MetBLUP: Phenotype prediction by selected metabolome

We next assessed whether predictability-based feature selection enhanced phenotype prediction. For metabolome-based models (MetBLUP; Fig. 4), models based on the most predictable features consistently outperformed those built from randomly selected subsets of equal size, and their predictive accuracy was often comparable to or greater than that of full-feature models.

**Figure 4.**
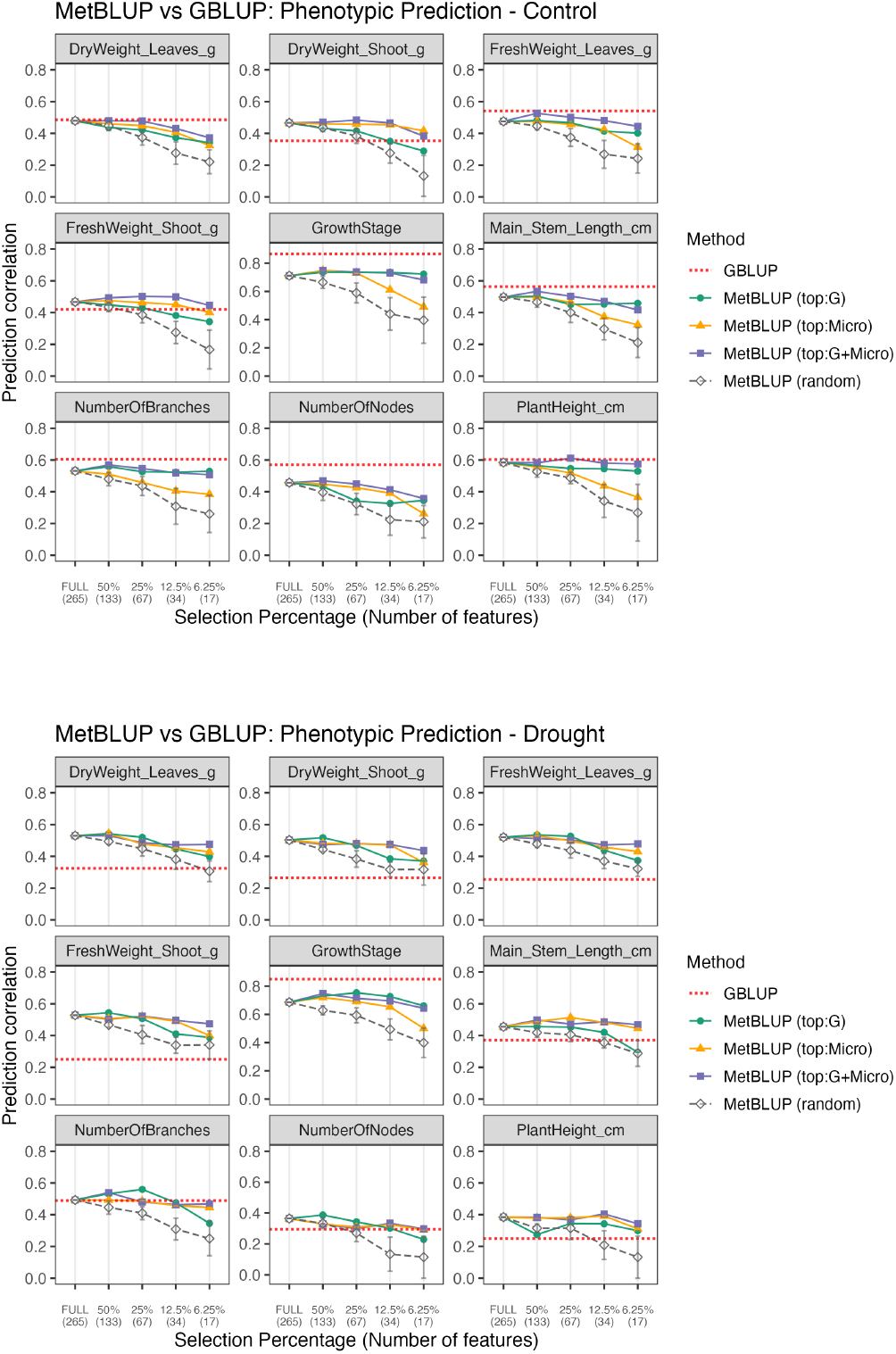
Phenotype prediction via MetBLUP across selection thresholds. The x-axis shows selection percentages, with the number of selected features at each threshold in parentheses. Solid lines represent models based on metabolome features selected by different methods: `“top:G”` (features selected using genomic information), `“top:Micro”` (features selected using microbiome information), and `“top:G+Micro”` (features jointly selected using both genomic and microbiome data). Dashed gray lines correspond to random feature selections matched by feature count. The dotted red horizontal lines indicate the prediction accuracy obtained by GBLUP, used here as a genomic baseline. For each point, the marker indicates the mean prediction correlation from 5-fold CV with five different seeds. For the `random` model, this procedure was repeated 10 times; the error bars represent the standard deviation across iterations. For the other models `(top:G, top:Micro, top:G+Micro)`, the kernel is identical across iterations; accordingly, no error bars are shown.

Overall, reducing the feature set from 265 to 34 features (25%) resulted in only a negligible reduction in prediction accuracy compared with using the full 265-feature set.

At each feature-count level, the predictability-based subsets consistently outperformed random subsets matched by feature count. Notably, the performance difference between the selected subsets and the random subsets became larger as the number of features decreased.

For most phenotypes, MetBLUP outperformed the genomic baseline (GBLUP) under drought conditions, and achieved comparable accuracy under control conditions.

When comparing the three feature-selection rules (top:G, top:Micro, and top:G+Micro), distinct trends emerged depending on the trait. The top:Micro performed best for traits with lower genomic predictability (e.g., shoot dry weight), whereas top:G yielded the highest accuracy for traits that were already well predicted by GBLUP (e.g., growth stage). The top:G+Micro rule provided complementary information, generally resulting in the most stable and robust performance across traits.

A similar trend was observed when comparing the two conditions, where GBLUP predictability was generally lower under drought, whereas MetBLUP based on top:G+Micro features achieved superior performance for major biomass traits (such as leaf dry weight), highlighting the influence of environmental factors mediated by metabolomic contributions.

#### 3.3.2. MicroBLUP: Phenotype prediction by selected microbiome

For microbiome-based models (MicroBLUP; Fig. 5), models using top predictable features also significantly outperformed random subsets of equal size. Under drought conditions, models using only the top 6.25% of microbiome features achieved predictive ability comparable to, or even exceeding, those using all features, and were on par with the GBLUP baseline for most biomass-related phenotypes, such as dry weight of leaves.

**Figure 5.**
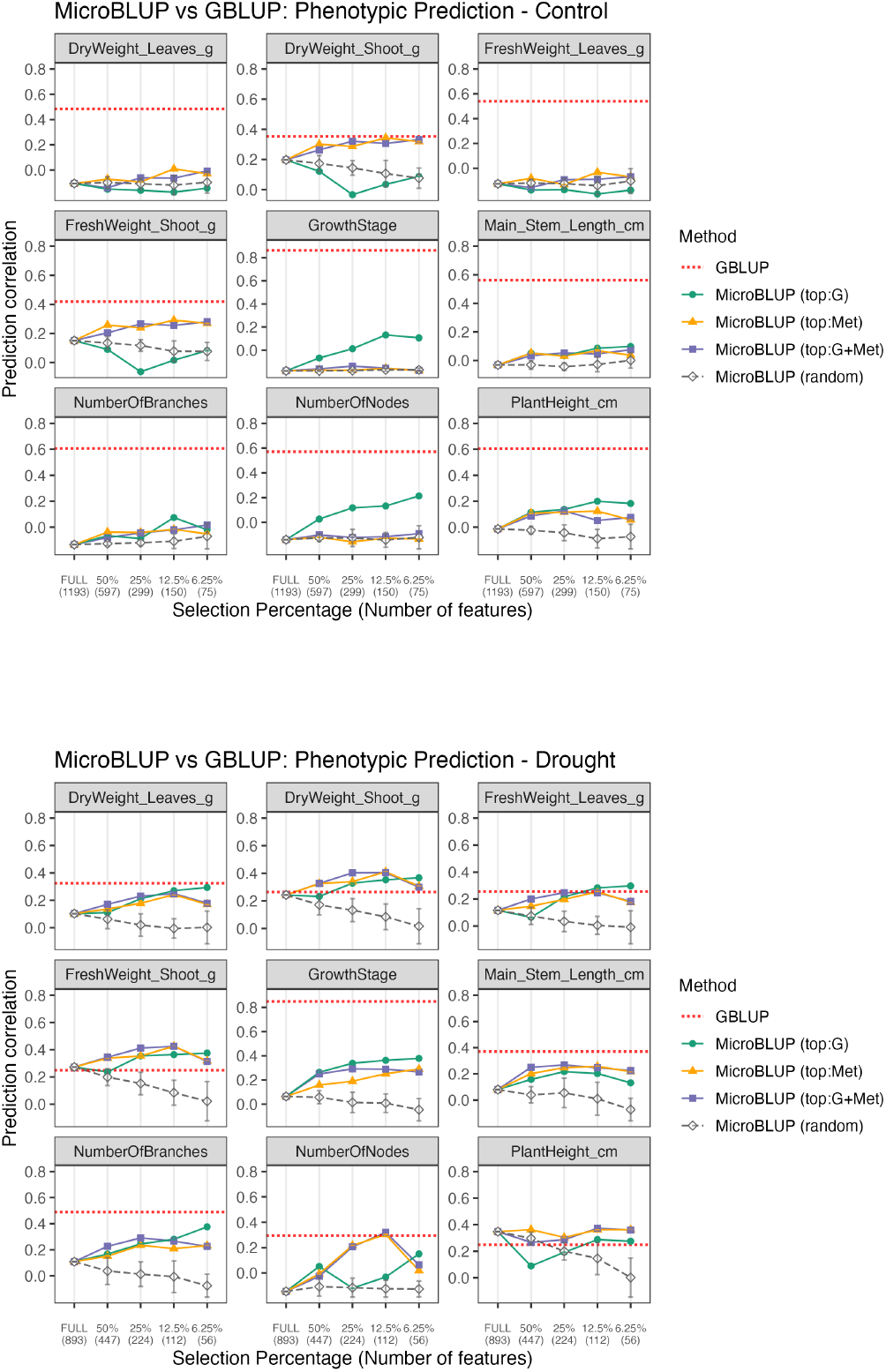
Phenotype prediction via MicroBLUP across selection thresholds. The x-axis shows selection percentages, with the number of selected features at each threshold in parentheses. Solid lines represent models based on microbiome features selected by different methods: `“top:G”` (features selected using genomic information), `“top:Met”` (features selected using metabolomic information), and `“top:G+Met”` (features jointly selected using both genomic and metabolomic data). Dashed gray lines correspond to random feature selections matched by feature count. The dotted red horizontal line indicates the prediction accuracy obtained by GBLUP, used here as a genomic baseline. For each point, the marker indicates the mean prediction correlation from 5-fold CV with 5 different seeds. For the `random` model, this procedure was repeated 10 times; the error bars represent the standard deviation across iterations. For the other models `(top:G, top:Met, top:G+Met)`, the kernel is identical across iterations; accordingly, no error bars are shown.

However, under control conditions, predictive ability remained generally lower than that of GBLUP. Notably, for major biomass-related traits such as shoot weight and leaf weight under control conditions, top:Met-based feature selection outperformed top:G, suggesting that microbiome features that are predictable from metabolomic data better capture relevant biological variation in these traits. In contrast, under drought conditions, the difference between the selection rules was smaller. In several cases where the microbiome was strongly filtered (6.25%), top:G achieved slightly higher accuracy than top:Met, indicating a shift in the dominant sources of microbiome predictability across environments.

## 4. Discussion

### 4.1. Predictive asymmetry between omics layers

Our results reveal a clear asymmetry in predictive power between the metabolome and microbiome layers. The metabolome was highly predictable from microbiome data, while the reverse direction showed weaker and more environment-sensitive patterns (Fig. 2). Moreover, the selected microbiome features were specific to each drought or control condition (Fig. 3). This finding is consistent with the notion that metabolite pools primarily represent downstream outputs of plant physiology, and that these metabolic outputs shape microbial community structure. As a result, the metabolome provides relatively stable biological signals, whereas the microbiome is more sensitive to environmental fluctuations.

### 4.2. Genetic versus environmental contributions to phenotype prediction

The predictability-based feature selection approach provides a unique perspective on how genetic and environmental factors shape phenotypic variation through different omics layers. Traits with high genomic predictability tended to show better phenotype prediction accuracy when using top:G-selected features, particularly in metabolome-based models (MetBLUP) (Fig. 4). This suggests that, for highly heritable traits, the metabolome captures genotype-associated physiological variation. In contrast, traits with lower genomic predictability or stronger environmental influence benefited more from top:Micro feature sets, indicating that metabolomic data also capture environmentally responsive biochemical or physiological processes that are not directly encoded in the genome.

### 4.3. Environmental modulation of omics–phenotype relationships

Under drought stress, genomic predictability (GBLUP accuracy) was generally reduced, whereas the phenotype prediction performance of MetBLUP and MicroBLUP models improved (Fig. 4, 5). This shift implies that environmental perturbation enhances the relevance of metabolomic and microbial responses to stress, leading to a greater contribution of environment-associated variation. Thus, while genetic control dominates under optimal conditions, environmental modulation of omic layers becomes a major driver of phenotypic variation under stress.

### 4.4. Hierarchical relationships among genome, metabolome, and microbiome

The relationships among the three omics layers imply a hierarchical structure. The genome and metabolome exhibited strong connectivity, consistent with a causal chain from genomic variation to metabolic state to phenotypic variation in biomass-related traits (Fig. 4). In contrast, the microbiome showed weaker direct associations with the genome, as evidenced by the poor performance of genome-based prediction of microbial composition, compared with metabolome-based prediction (Fig. 3). We also observed the limited ability of the top:G-based MicroBLUP model to predict phenotypic traits, particularly under control conditions (Fig. 5). In addition, microbial composition was explained substantially better by metabolomic data than by genomic data. Given that metabolomic features were comparably predictable from both the genome and the microbiome (Fig. 3), this pattern suggests a potential hierarchical structure in which genomic variation shapes metabolomic states, which in turn influence rhizosphere microbial communities.

### 4.5. Biomarker selection

Our reciprocal framework provides a practical approach to prioritizing omics features based on their predictability and leveraging them for phenotype prediction. In the metabolome, the filtered feature sets (25%) performed comparably to the full dataset in the MetBLUP framework, whereas in the microbiome, an even smaller subset of features (6.25%) outperformed the full dataset in the MicroBLUP framework (Fig. 4, 5). These selected subsets highlight candidate biomarkers in both metabolomic and microbiome layers, improving biological interpretability. For the metabolome in particular, reducing the number of features has practical benefits, as targeted assays can become substantially less costly when only a small set of compounds needs to be measured.

## 5. Conclusions

Our findings highlight that integrating predictability-based feature selection enables the disentanglement of genetic and environmental contributions to complex traits. The proposed framework clarifies the distinct yet interconnected roles of the genome, metabolome, and microbiome layers. Moreover, our approach enables the selection of biologically meaningful features and enhances the performance of phenotype prediction models. Such an understanding is critical for improving multi-omics prediction models and for designing strategies to exploit both genetic and environmental variation in crop improvement.

## Author Contributions

Conceptualization, H.Y., G.M. and H.I.; Data curation, Y.F., and Y.I.; Formal analysis, H.Y.; Supervision, H.I.; Writing – original draft, H.Y.; Writing – review & editing, G.M., H.I.

## Data Availability Statement

All source codes and data are available from the repository in GitHub: https://github.com/Yoska393/ReciprocalBLUP.

## Acknowledgments

We are grateful to Yushiro Fuji, Yui Nose, and Yasunori Ichihashi at RIKEN for their support in data collection. We also thank the technical staff at the Arid Land Research Center, Tottori University, for their assistance.

## Appendix A

### Multi-omics data collection

#### Appendix A.1 Field experiment

A total of 198 diverse soybean accessions from the NARO Genebank (https://www.gene.affrc.go.jp/), representing the Global Soybean Minicore Collection [4,24], were cultivated at the Arid Land Research Center of Tottori University in 2019. Field management followed Dang et al. [17], Toda et al. [25], Sakurai et al. [26]. Plants were grown on sandy soil under two watering regimes: a well-watered control and a drought (non-irrigated) condition. Fertilizers were applied prior to sowing, and irrigation was controlled through buried drip tubes under white mulch sheets. Biomass-related phenotypic measurements and rhizosphere root samples were collected from the same plots at the beginning of September to ensure data consistency across omics layers. Root samples were immediately frozen at *−*80 ^*°*^*C* for metabolome and microbiome analyses. The soybean genome sequences are available from the DDBJ Sequence Read Archive (BioProject accession: PRJDB7281).

#### Appendix A.2 Phenotypic traits

Nine biomass-related traits were measured: dry and fresh weights of shoots and leaves, growth stage, plant height, stem length, number of nodes, and number of branches.

#### Appendix A.3 Metabolome analysis

Root metabolomic profiling was conducted following Sawada et al. [27] and Uchida et al. [28]. Freeze-dried and powdered root samples were extracted using 80% methanol containing internal standards, analyzed using LC–QqQ–MS (LCMS-8050, Shimadzu, Kyoto, Japan), and quantified in multiple reaction monitoring (MRM) mode. Peak areas were processed using MRMPROBS [29] and normalized against internal standards. The processed metabolomic dataset is available from the RIKEN DropMet database (ID: DM0071).

#### Appendix A.4 Microbiome analysis

Root-associated and rhizosphere bacterial communities were characterized using 16S rRNA gene sequencing. Genomic DNA was extracted following Kumaishi et al. [30] and Ichihashi et al. [31], and the V4 region of the 16S rRNA gene was amplified using a two-step PCR protocol with PNA blocking primers [32,33]. Amplicons were sequenced on an Illumina MiSeq platform (2*×*300 bp). Raw reads were processed using QIIME2 (ver. 2020.6.0) with Cutadapt and DADA2 [34] for quality filtering, denoising, and chimera removal. Taxonomic assignment was performed with the SILVA database (ver. 138) [35,36], and non-bacterial sequences were excluded. Rarefaction was performed (*n* = 3000), resulting in 1,642 ASVs. Additional filtering was applied separately for each condition. The final dimensions of the data were 179 *×* 1193 for the control condition and 182 *×* 893 for the drought condition. Raw sequencing data are available at DDBJ (BioProject accession: PRJDB37881).

### Disclaimer/Publisher’s Note

The statements, opinions and data contained in all publications are solely those of the individual author(s) and contributor(s) and not of MDPI and/or the editor(s). MDPI and/or the editor(s) disclaim responsibility for any injury to people or property resulting from any ideas, methods, instructions or products referred to in the content.

